# Giant viruses encode novel types of actins possibly related to the origin of eukaryotic actin: the viractins

**DOI:** 10.1101/2020.06.16.150565

**Authors:** Violette Da Cunha, Morgan Gaia, Hiroyuki Ogata, Olivier Jaillon, Tom O. Delmont, Patrick Forterre

## Abstract

Actin is a major component of the eukaryotic cytoskeleton. Many related actin homologues can be found in eukaryotes1, some of them being present in most or all eukaryotic lineages. The gene repertoire of the Last Eukaryotic Common Ancestor (LECA) therefore would have harbored both actin and various actin-related proteins (ARPs). A current hypothesis is that the different ARPs originated by gene duplication in the proto-eukaryotic lineage from an actin gene that was inherited from Asgard archaea. Here, we report the first detection of actin-related genes in viruses (viractins), encoded by 19 genomes belonging to the Imitervirales, a viral order encompassing the giant Mimiviridae. Most viractins were closely related to the actin, contrasting with actin-related genes of Asgard archaea and Bathyarchaea (a newly discovered clade). Our phylogenetic analysis suggests viractins could have been acquired from proto-eukaryotes and possibly gave rise to the conventional eukaryotic actin after being reintroduced into the pre-LECA eukaryotic lineage.

## Introduction

Actin is a major component of the eukaryotic cytoskeleton. Many related actin homologues can be found in eukaryotes^1^, some of them being present in most or all eukaryotic lineages^2^. The gene repertoire of the Last Eukaryotic Common Ancestor (LECA) therefore would have harbored both actin and various actin-related proteins (ARPs)^1,2^. A current hypothesis is that the different ARPs originated by gene duplication in the proto-eukaryotic lineage from an actin gene that was inherited from Asgard archaea^3,4^. Here, we report the first detection of actin-related genes in viruses (viractins), encoded by 19 genomes belonging to the *Imitervirales*, a viral order encompassing the giant *Mimiviridae*^5^. Most viractins were closely related to the actin, contrasting with actin-related genes of Asgard archaea and *Bathyarchaea* (a newly discovered clade). Our phylogenetic analysis suggests viractins could have been acquired from proto-eukaryotes and possibly gave rise to the conventional eukaryotic actin after being reintroduced into the pre-LECA eukaryotic lineage.

### The discovery of viractins

We first detected an actin-like gene (thereafter dubbed viractin) in the giant virus *Yasminevirus*, recently isolated from sewage water by means of amoeba coculture^6^. *Yasminevirus* belongs to the *Mimiviridae* family^7,8^ within the proposed *Klosneuvirinae*^6,9^ subfamily. *Mimiviridae* are giant viruses belonging to the *Nucleocytoviricota*^5^ viral phylum, previously known as the NucleoCytoplasmic Large DNA virus assemblage (NCLDV)^7^. Using the *Yasminevirus* viractin gene as a query, we detected additional viractins in 16 metagenome-assembled genomes (MAGs) of *Nucleocytoviricota* originating mostly from marine and freshwater systems^9–11^, as well as in two additional MAGs of *Nucleocytoviricota* we characterized from the sunlit ocean (see method section). Table 1 summarizes genomic statistics for the NCLDV isolate genome and MAGs containing a viractin. MAGs were affiliated to Hokovirus (n=1) and *Yasminevirus* (n=1) within *Klosneuvirinae*, as well as to two lineages related to *Mimiviridae* dubbed MVGL55 (n=15) and MM15 (n=1) that were recently characterized from large metagenomic surveys^10,11^. These clades correspond to the newly revealed diversity of *Mimiviridae* relatives that have been recently included into the *Imitervirales*^5^ order. The position of the viruses encoding viractin within the *Imitervirales* order was confirmed with a phylogenetic reconstruction of representative sequences using previously studied markers and datasets (Fig S1). Notably, at least two *Imitervirales* lineages (Yasminevirus-like and MVGL55) are enriched in viractin, suggesting a specific recruitment of actin and actin-related proteins by the viral common ancestors of these clades instead of recent and independent multiple acquisitions in few *Mimiviridae-*related genomes. The actin encoding MVGL55 viruses were identified in lakes (Lanier and Michigan Lake), oceans (Atlantic ocean, Pacific ocean, Arctic ocean), and seas (North sea and Mediterranean sea)(Table 1, table S1). Moreover, the two Yasminevirus encoding viractins were characterized from very different environments, sewage water from Jeddah in Saudi Arabia and the Pacific Ocean. Thus, while the isolation of Yasminevirus provides proof of the existence of viractin, environmental surveys (including the metagenomic harvest of *Tara* Oceans expeditions^12^) reveal that not one but multiple clades of *Mimiviridae*-related viruses, within and beyond the *Klosneuvirinae* encode viractin, always found in single copy.

**Table 1:** Summary of 19 genomes of *Imitervirales* containing a viractin. Statistics were processed with anvi’o13. Completion was estimated using HMMs targeting eight NCLDV gene markers8. The identity is given for a coverage ranging from 99% to 100% to the human actin (* for viractin 04 the identity is given for a coverage of 73%). Details in table S1.

| NCLDV families containing a viractin | Clade of genomes | Source | Size | GC content | Estimated completion | Ecosystem | Viractin lineage | Identity to human actin |
| --- | --- | --- | --- | --- | --- | --- | --- | --- |
| Mimiviridae-related | Clade MVGL55 MAG | Schultz et al. 2020 | 0.40 Mbp | 37.9% | 87.5% | Freshwater | Viractin 01 | 71.8% |
|  | Clade MVGL55 MAG | Schultz et al. 2020 | 0.40 Mbp | 36.8% | 100% | Freshwater |  | 73.6% |
|  | Clade MVGL55 MAG | Schultz et al. 2020 | 0.29 Mbp | 36.5% | 62.5% | Freshwater |  | 73.4% |
|  | Clade MVGL55 MAG | Moniruzzaman et al. 2020 | 0.38 Mbp | 34.1% | 87.5% | Marine |  | 71.5% |
|  | Clade MVGL55 MAG | Schultz et al. 2020 | 0.25 Mbp | 32.0% | 100% | Marine |  | 67.3% |
|  | Clade MVGL55 MAG | Schultz et al. 2020 | 0.44 Mbp | 29.9% | 100% | Marine |  | 71.4% |
|  | Clade MVGL55 MAG | Schultz et al. 2020 | 0.46 Mbp | 33.0% | 100% | Marine |  | 69.2% |
|  | Clade MVGL55 MAG | Schultz et al. 2020 | 0.35 Mbp | 29.7% | 87.5% | Marine |  | 68.6% |
|  | Clade MVGL55 MAG | Schultz et al. 2020 | 0.41 Mbp | 32.1% | 87.5% | Marine |  | 71.5% |
|  | Clade MVGL55 MAG | Schultz et al. 2020 | 0.32 Mbp | 34.3% | 87.5% | Marine |  | 64.7% |
|  | Clade MVGL55 MAG | Schultz et al. 2020 | 0.52 Mbp | 34.5% | 100% | Marine |  | 71.5% |
|  | Clade MVGL55 MAG | Schultz et al. 2020 | 0.38 Mbp | 35.5% | 75% | Marine |  | 71.2% |
|  | Clade MVGL55 MAG | Schultz et al. 2020 | 0.29 Mbp | 35.9% | 50% | Marine |  | 69% |
|  | Clade MVGL55 MAG | Schultz et al. 2020 | 0.33 Mbp | 31.6% | 100% | Marine |  | 71% |
|  | Clade MVGL55 MAG | TARA Oceans consortium | 0.28 Mbp | 34.6% | 87.5% | Marine |  | 72% |
| Mimiviridae-related | Clade MM15 MAG | Moniruzzaman et al. 2020 | 0.71 Mbp | 24.1% | 87.5% | Marine | Viractin 02 | 66.9% |
| Mimiviridae (Klosneuvirinae sub-family) | Yasminevirus isolate | Bajrai et al. 2019 | 2.13 Mbp | 40.3% | 100% | Sewage | Viractin 03 | 59.2% |
|  | Yasminevirus MAG | TARA Oceans consortium | 1.45 Mbp | 45.9% | 100% | Marine |  | 55.4% |
| Mimiviridae (Klosneuvirinae sub-family) | Hokovirus MAG | Schultz et al. 2017 | 1.33 Mbp | 21.4% | 100% | Wastewater | Viractin 04 | 27% * |

### At least four lineages of viractins in the *Imitervirales* order

Since viractin was previously unknown in the viral world, we hypothesized that *Imitervirales* containing viractin recruited this gene from their hosts. We performed a phylogenetic analysis to determine if this recruitment occurred only once or several times independently, and to possibly find out the original eukaryotic host(s). We included the 19 newly discovered viractins together with actin sequences, and those of various clades of eukaryotic ARPs (ARP-01 – also called centractin - to ARP-10). We also included all ARP sequences recently discovered in Asgard archaea (hereafter asgardactins) and a new group of ARPs, hereafter dubbed as bathyactins, that we unexpectedly identified in some *Bathyarchaea*^14,15^ (see table S1). It is, to our knowledge, the first time that such a closely related homologue of eukaryotic actin is detected in Archaea not belonging to the Asgard archaea. These *Bathyarchaea* additionally encode crenactin, a more distantly related actin-like protein encoded by most *Crenarchaea, Bathyarchaea, Aigarchaea* and *Korarchaea* (Fig S2; see Method for the selection of sequences). We used these archaeal crenactins as the outgroup for rooting (Fig 1, Fig S2).

**Figure 1:**
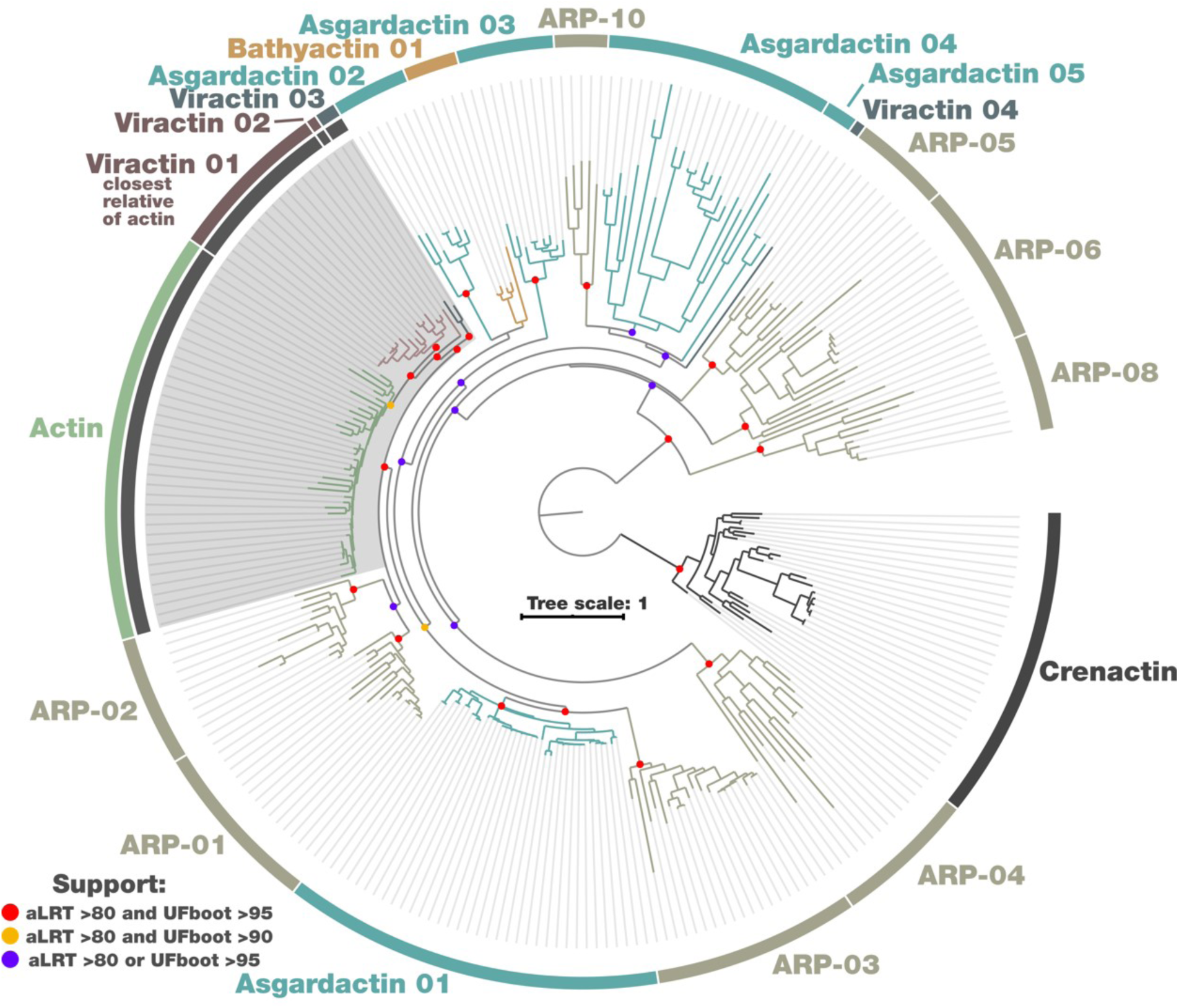
Phylogenetic tree of the EL-actin clade rooted with crenactin. The EL-actin clade includes eukaryotic actin and ARPs, asgardactins, and newly identified bathyactins and viractins. The scale-bar indicates the average number of substitutions per site. Detailed tree in Fig S2.

In our phylogenetic tree (Fig 1), no sequence nor clade was branching close to the crenactin outgroup, suggesting that bathyactins, asgardactins, and viractins are indeed all related to the eukaryotic actins and ARPs, forming a large clade hereafter dubbed the EL-actin (Eukaryotic-Like actin) clade. The actin and all eukaryotic ARP formed individual monophyletic clades that were clearly separated from each other in our tree (Fig 1). The 19 viractins were structured in four clades, viractin 01-04, corresponding to the four different groups of *Imitervirales* encoding viractins (see Table 1, Fig 1). On the front of Archaea, all bathyactins were grouped in a single clade while the asgardactins were separated in five clades (herein called asgardactin 01-05), in line with recent phylogenetic analyses^16^. Notably, the five clades of asgardactins and the clade of bathyactin were not located at the base of the EL-actin clade, as expected if eukaryotic actin and ARPs originated from these archaeal actins, but branched at different positions between different clades of eukaryotic ARPs. Two clades of asgardactins only include one phylum of Asgardarchaea (*Lokiarchaeota* for asgardactin 02, 03), and only asgardactin 01 is present in all phyla of Asgard archaea. As in the case of viractins, this suggests multiple recruitments of ancestral actin-related proteins from proto-eukaryotes by these Archaea, similarly to a described case of horizontal gene transfer (HGT) of actin between eukaryote and bacteria^19^. Comparing the predicted structure models of one reference sequence for each of the different clades of viractin, asgardactin, bathyactin, and ARPs proteins with the actin structure revealed that all predicted structural domains are conserved in EL-actins, indicating that these proteins could share similar biochemical functions (Fig S2).

### The origin of viractins

Surprisingly, viractins did not branch within either the eukaryotic actins or any of the eukaryotic clades of ARPs, as would be expected in the case of a recent transfer from modern eukaryotes to *Imitervirales*. Instead, these monophyletic viractin clades branched at two positions between the different eukaryotic clades (Fig 1). Viractins 01, 02, and 03 were basal to the actin, whereas the shorter viractin 04 (ca. 75% of the average length) branched at the root of a clade grouping ARP-10 and asgardactin 04-05, clearly indicating that viractins were recruited at least twice independently by different *Mimiviridae-*related clades.

Importantly, the eukaryotic actin and all ARP clades but ARP-10 include protists and pluricellular eukaryotes from different supergroups, including Amorphea, Archaeplastida, TSAR and Excavates^17^, indicating that they were most likely acquired before the emergence of modern eukaryotes and were hence already present in LECA. Consequently, most nodes at the base of each of these eukaryotic clades correspond to the relative position of LECA (Fig S2). The basal position of the viractins could be an indication that viractins evolved more rapidly than ARPs and actin and were artificially attracted in our phylogenetic reconstruction outside of the cellular clades they should be branching with by a phenomenon of long branch attraction (LBA), however the lengths of their branches were similar to those of their cellular counterparts. It hence seems more likely that they were recruited by ancient *Imitervirales* from proto-eukaryotes, before LECA and the diversification of modern eukaryotes. The topology of the phylogenetic tree with viractins 01 to 03, corresponding to different *Mimiviridae*-related lineages (Table 1 and Fig S1), closely related to each other but not as a single monophyletic clade, suggests a complex evolutionary history of transfers and losses through the co-evolution of *Imitervirales* and their hosts. Interestingly, this topology implies that the *Nucleocytoviricota* were not only already diversified at the family level before LECA, as suggested previously from analyses on the DNA-dependent RNA polymerase^8^ and taxon richness and diversity^18^, but also at the subfamily level for the *Mimiviridae*.

Finally, the intriguing positions of actin and viractins 01, 02, and 03 (grey area) suggests that actin might have a viral origin, with first an actin-related gene captured by a specific *Mimiviridae-*related clade where it evolved before being transferred back to the pre-LECA-eukaryotic lineage. The discovery of these viractin lineages (and to a lesser extent, the bathyactins) provides a new perspective regarding the evolutionary history of the eukaryotic cytoskeleton but stresses the need for extensive phylogenetic studies of the newly extended EL-actin clade.

## Discussion

Actin and ARPs are paramount features of the eukaryotic cytoskeleton involved in various cellular processes that were already present in LECA^2^. Here, we report the first identification of actin-like genes in the viral world, which we dubbed viractins. We identified single-copy viractins in four different clades of viruses related to the *Mimiviridae* family in the *Imitervirales* order. Our results point at a diversification of *Imitervirales* before the emergence of modern eukaryotes. The discovery of three viractin clades closely related to the eukaryotic actin echoes the close association between *Imitervirales* and several eukaryotic signature features (DNA-dependent RNA polymerase II^8^, histones^20^ and DNA polymerase^21^), substantiating the hypothesis of a co-evolution between NCLDVs and proto-eukaryotes that played a major contribution in shaping the molecular components and functions of the modern eukaryotic cell. Additional viractin sequences and an improved understanding of the phylogeny of eukaryotes are now needed to refine the model of evolution regarding the origin of actin.

Different authors previously emphasized a possible evolutionary relationship between the eukaryotic nucleus and NCLDV viral factories^22,23^. In this context, it is interesting to note that actin is present in both the cytoplasm and nucleus of eukaryotic cells and that, in collaboration with nuclear ARPs, seems to be involved in several nuclear-related processes^24^. The viral eukaryogenesis hypotheses, which attribute various roles in the emergence of eukaryotes to viruses, have notably been recently boosted by the discovery of Caudovirales lineages producing a nucleus-like structure within infected bacterial cells^25^. These viruses encode a distant homologue of the eukaryotic tubulin that localizes this nucleus-like structure in the middle of the virocell (i.e. the virus-infected cell) and treadmills toward it the viral capsids that were assembled on the membrane^26^. It is possible that viractins play a similar role during viral infections by controlling the localization of the viral factory close to the host nucleus (e.g., as seen in *Yasminevirus*^6^). Deciphering the role of viractins during viral infection will be an exciting challenge for the future.

## Method

### Viractin identification

The first viractin protein (VBB18706, 377 aa) was detected in the giant virus Yasminevirus by screening viruses in the NCBI nr database by BLAST search using *Homo sapiens* actin cytoplasmic 1 (NP_001092.1, 375 aa) as query. In order to identify potential new viractin genes in giant viruses, we searched for additional viractins in NCLDV MAGs originating mostly from marine and freshwater systems^9,10,27^, using one HMM dedicated to the identification of actin. In screening for actin-related proteins in Archaea, we identified by BLASTP a new clade of ARPs in some *Bathyarchae*a, and we also extracted ARP encoded by the Asgard archaea^3,28–31^.

### Genome-resolved metagenomics

Two metagenome-assembled genomes (MAGs) of *Mimiviridae* containing a viractin were characterized from metagenomes of TARA Oceans (Pacific Ocean and Mediterranean Sea), by performing manual binning and curation on large size fractions of surface ocean plankton (0.8-2,000 microns)^32^. Briefly, we used the anvi’o platform^33^ and a co-assembly strategy followed by binning using sequence composition and differential coverage, as previously applied to a small size fraction of surface ocean plankton (0.2-3 microns)^34^. We searched for NCLDV MAGs containing a viractin using eight HMMs of gene markers for NCLDV^8^ and the HMM dedicated to identification of actin.

### *Nucleocytoviricota* phylogenetic tree and *Imitervirales* order schematic tree

The phylogenetic tree of *Nucleocytoviricota* in Fig S1 was performed on the concatenation of the two largest DNA-dependent RNA polymerase subunits with the protocol and dataset detailed in^8^: the model was estimated using ModelFinder Plus option in IQ-TREE version 1.6.12, and supports were computed from 1,000 replicates for the Shimodaira-Hasegawa (SH)-like approximation likelihood ratio test (aLRT)^35^ and ultrafast bootstrap approximation (UFBoot)^36^. The schematic tree displayed in Fig S1 was made starting from a reference phylogenetic tree of various NCLDV clades^11^ (see https://doi.org/10.6084/m9.figshare.11774958.v1). We pruned the part of the tree corresponding to *Mimiviridae* and incorporated the position of Yasminevirus^6^.

### Actin superfamily phylogeny

To position the viractin within the actin superfamily, we extracted representative sequences using reference datasets^1,2^, and incorporated other actins and ARPs to expend the dataset. Finally, we added archaeal sequences corresponding to crenactin as an outgroup. The alignment was performed using MAFFT v7.45^37^. Aligned sequences were trimmed with BMGE, with the -m BLOSUM 30 and -b1 options^38^. The maximum likelihood trees were constructed using IQ-TREE version 1.6.12^39^ under the LG+R5 model of evolution according the MFP option for model selection. The branches support was calculated by the SH-like aLRT (10,000 replicates) and UFBoot (10,000 replicates)^35,36^.

### Visualization

Visualization of the phylogenetic tree in Figs 1 was performed using the anvi’o interactive interface set in manual mode. Tree scale was incorporated using iTOL^41^, and support information was manually added in Inkscape. The other trees were visualized using iTOL.

### Structure prediction

Structure prediction of a representative of each group of viractin (Yasminevirus, MM15, MM1, Hokovirus), asgardactin (*candidatus* Prometheoarchaeum syntrophicum QEE15133,17499,17026,15652), and *candidatus* Heimdallarchaeota archaeon (RLI71419), and bathyactin (RLI09076) was made using Phyre2^42^. The obtained predicted structures were compared to the structure of the uncomplexed actin (1J6Z) of *Oryctolagus cuniculus*.

### Data availability

All data our study used was made publically available. This includes (1) the HMM to search for actin related genes in genomes and metagenomic assemblies (https://figshare.com/s/0cbbc12cce5bce2ed3a1), (2) the anvi’o summary of 19 NCLDV genomes containing a viractin (https://figshare.com/s/23a8419492b3ae726f39), (3) raw phylogenetic trees for figures 1 and 2 (https://figshare.com/s/e40401fc13331992730a), (4) the FASTA sequences of amino acid sequences used in the EL-actin phylogenetic analyses (https://figshare.com/s/c3af4f2cdce6b3bb5a11), (5) and data related to predicted protein structures (https://figshare.com/s/052304ec83f13ed41411).

---

**Supplementary Figure 1.**
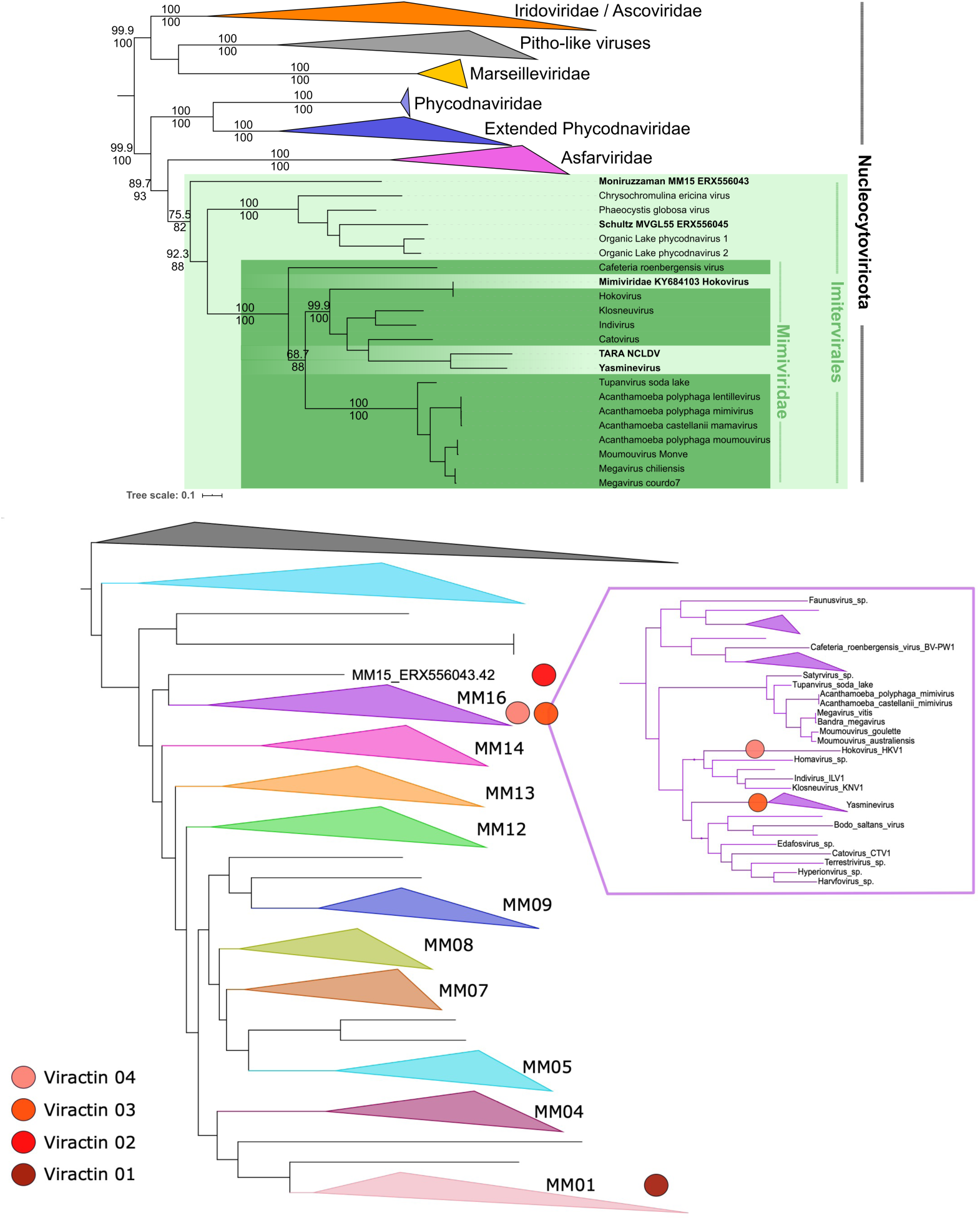
Upper panel: phylogenetic tree of the *Nucleocytoviricota* with representatives of the four clades of viractins, using the concatenation of the two largest DNA-dependent RNA polymerase subunits. Values at nodes computed by aLRT and UFBoot. The scale-bar indicates the average number of substitutions per site. Lower panel: schematic phylogeny of *Imitervirales*. This schema was made based on the NCLDV phylogenetic tree11 and we added the position of the Yasminevirus6.

**Supplementary Figure 2.**
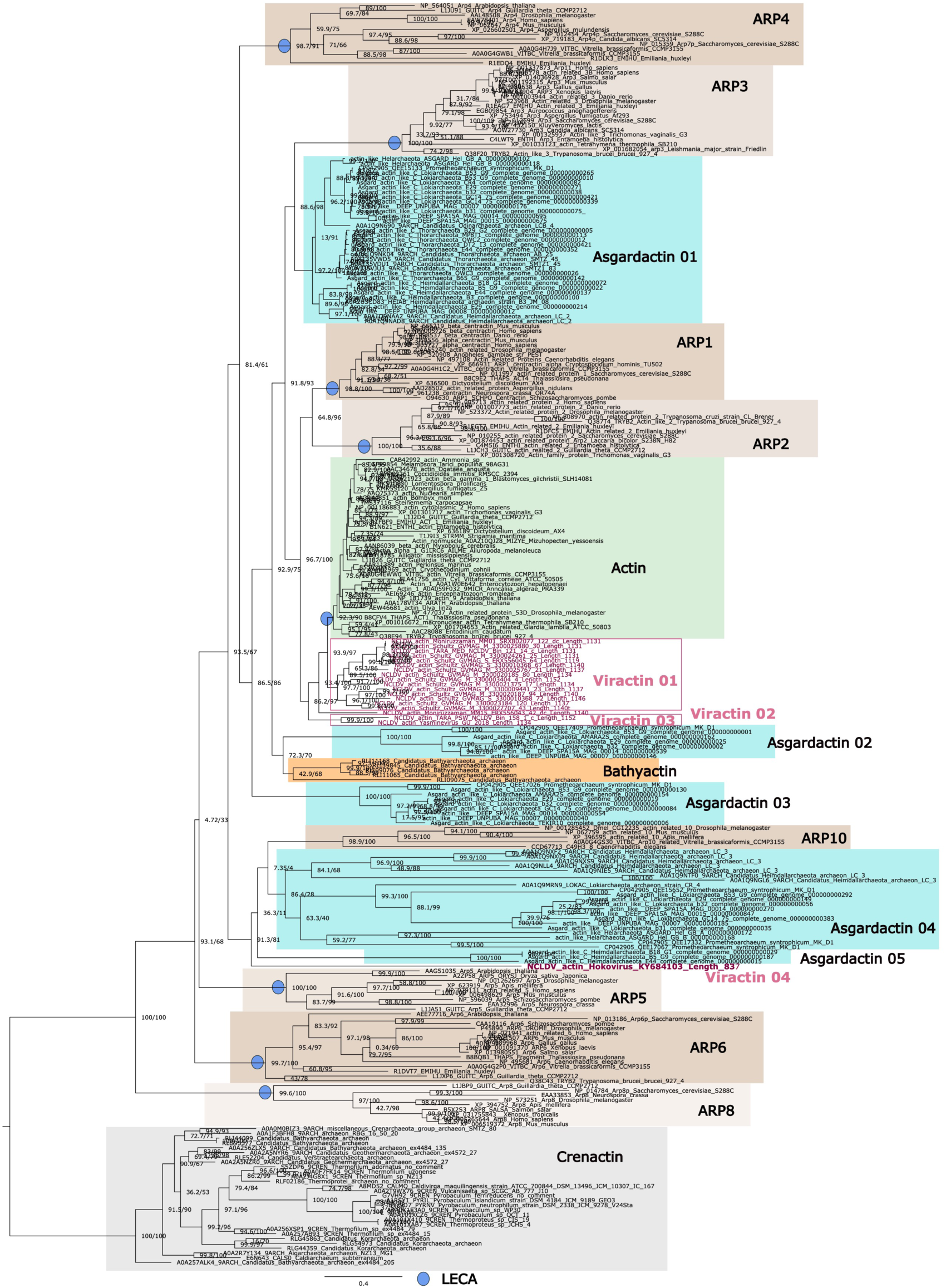
Maximum Likelihood (ML) phylogenetic tree of actin and actin-related proteins (296 positions, model LG+R5) rooted with the crenactin. The eukaryotic actin and ARPs are indicated in green and brown, respectively. The viractins are indicated in pink. The asgardactin and the bathyactin are indicated in blue and orange, respectively. The multiple position of LECA on the tree are indicated with blue dots. Values at nodes represent support calculated with the SH-like aLRT (10,000 replicates) and UFBoot (10,000 replicates). The scale-bars represent the average number of substitutions per site.

**Supplementary Figure 3.**
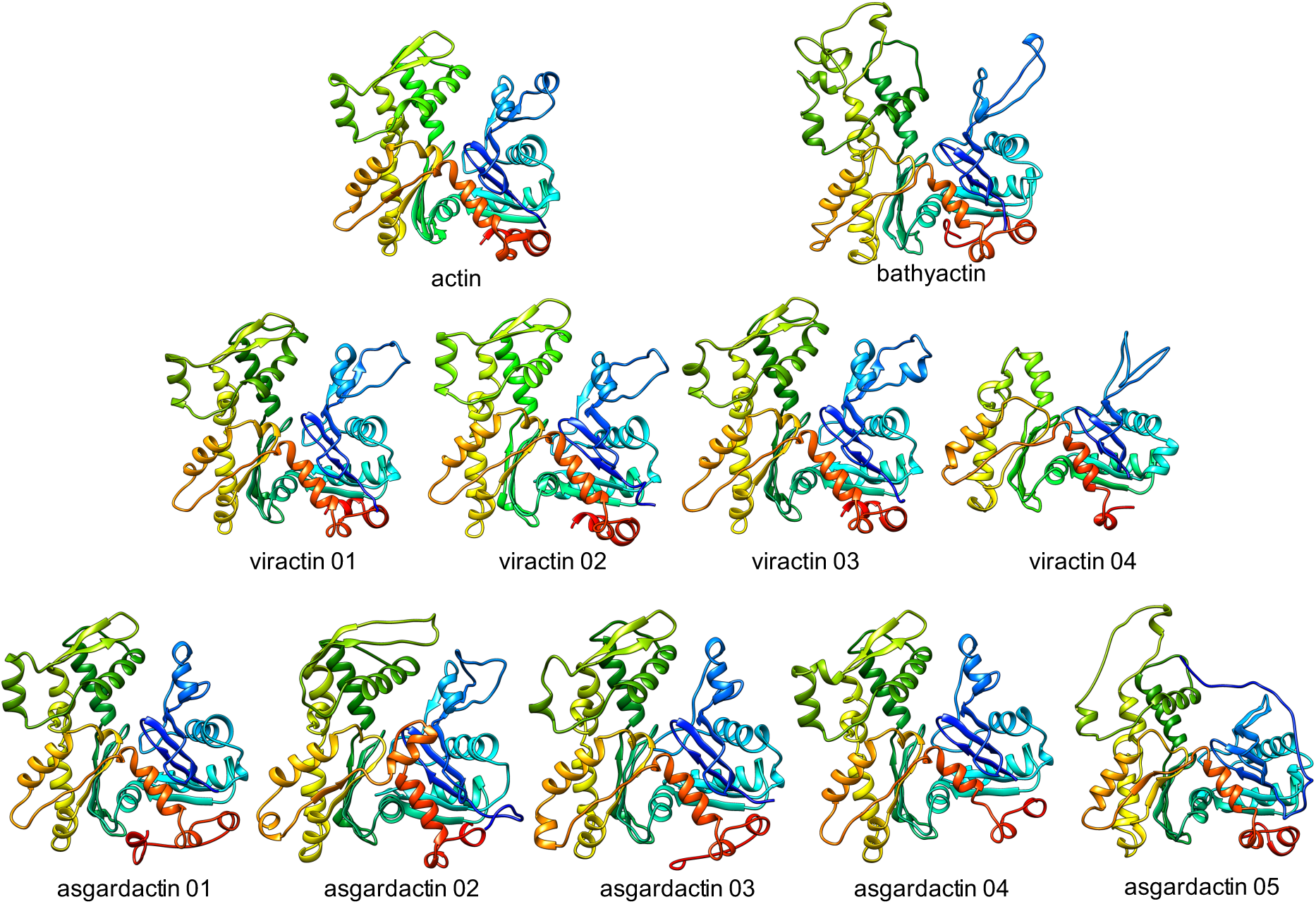
Comparison of the uncomplexed actin structure to the predicted structures of a reference sequences of viractins, asgardactins and bathyactin clades.

## Notes

### Competing Interest Statement

The authors have declared no competing interest.

## Bibliography

1. Goodson, H. V. & Hawse, W. F. Molecular evolution of the actin family. J. Cell Sci. (2002).

2. Muller, J. et al. Sequence and comparative genomic analysis of actin-related proteins. Mol. Biol. Cell (2005). doi: 10.1091/mbc.E05-06-0508

3. Zaremba-Niedzwiedzka, K. et al. Asgard archaea illuminate the origin of eukaryotic cellular complexity. Nature 541, 353–358 (2017).

4. Lindås, A. C., Valegård, K. & Ettema, T. J. G. Archaeal actin-family filament systems. Subcell. Biochem. (2017). doi: 10.1007/978-3-319-53047-5_13

5. Koonin, E. V. et al. Global Organization and Proposed Megataxonomy of the Virus World. Microbiol. Mol. Biol. Rev. (2020). doi: 10.1128/mmbr.00061-19

6. Bajrai, L. H. et al. Isolation of Yasminevirus, the First Member of Klosneuvirinae Isolated in Coculture with Vermamoeba vermiformis, Demonstrates an Extended Arsenal of Translational Apparatus Components. J. Virol. (2019). doi: 10.1128/jvi.01534-19

7. Koonin, E. V. & Yutin, N. Evolution of the Large Nucleocytoplasmic DNA Viruses of Eukaryotes and Convergent Origins of Viral Gigantism. in Advances in Virus Research (2019). doi: 10.1016/bs.aivir.2018.09.002

8. Guglielmini, J., Woo, A. C., Krupovic, M., Forterre, P. & Gaia, M. Diversification of giant and large eukaryotic dsDNA viruses predated the origin of modern eukaryotes. Proc. Natl. Acad. Sci. (2019). doi: 10.1073/pnas.1912006116

9. Schulz, F. et al. Giant viruses with an expanded complement of translation system components. Science (80-.). (2017). doi: 10.1126/science.aal4657

10. Schulz, F. et al. Giant virus diversity and host interactions through global metagenomics. Nature (2020). doi: 10.1038/s41586-020-1957-x

11. Moniruzzaman, M., Martinez-Gutierrez, C. A., Weinheimer, A. R. & Aylward, F. O. Dynamic genome evolution and complex virocell metabolism of globally-distributed giant viruses. Nat. Commun. (2020). doi: 10.1038/s41467-020-15507-2

12. Bork, P. et al. Tara Oceans studies plankton at planetary scale. Science (80-.). 348, 873 (2015).

13. Eren, A. M. et al. Anvi’o: an advanced analysis and visualization platform for ‘omics data. PeerJ 3, e1319 (2015).

14. Zhou, Z., Pan, J., Wang, F., Gu, J. D. & Li, M. Bathyarchaeota: Globally distributed metabolic generalists in anoxic environments. FEMS Microbiology Reviews (2018). doi: 10.1093/femsre/fuy023

15. Dombrowski, N., Teske, A. P. & Baker, B. J. Expansive microbial metabolic versatility and biodiversity in dynamic Guaymas Basin hydrothermal sediments. Nat. Commun. (2018). doi: 10.1038/s41467-018-07418-0

16. Stairs, C. W. & Ettema, T. J. G. The Archaeal Roots of the Eukaryotic Dynamic Actin Cytoskeleton. Current Biology (2020). doi: 10.1016/j.cub.2020.02.074

17. Burki, F., Roger, A. J., Brown, M. W. & Simpson, A. G. B. The New Tree of Eukaryotes. Trends in Ecology and Evolution (2020). doi: 10.1016/j.tree.2019.08.008

18. Mihara, T. et al. Taxon richness of “Megaviridae” exceeds those of bacteria and archaea in the ocean. Microbes Environ. (2018). doi: 10.1264/jsme2.ME17203

19. Guljamow, A., Delissen, F., Baumann, O., Thünemann, A. F. & Dittmann, E. Unique properties of eukaryote-type actin and profilin horizontally transferred to cyanobacteria. PLoS One (2012). doi: 10.1371/journal.pone.0029926

20. Yoshikawa, G. et al. Medusavirus, a Novel Large DNA Virus Discovered from Hot Spring Water. J. Virol. (2019). doi: 10.1128/jvi.02130-18

21. Takemura, M., Yokobori, S. I. & Ogata, H. Evolution of Eukaryotic DNA Polymerases via Interaction Between Cells and Large DNA Viruses. J. Mol. Evol. (2015). doi: 10.1007/s00239-015-9690-z

22. Forterre, P. & Gaïa, M. Giant viruses and the origin of modern eukaryotes. Current Opinion in Microbiology (2016). doi: 10.1016/j.mib.2016.02.001

23. Bell, P. J. Evidence supporting a viral origin of the eukaryotic nucleus. bioRxiv (2019). doi: 10.1101/679175

24. Bajusz, C. et al. Nuclear actin: ancient clue to evolution in eukaryotes? Histochemistry and Cell Biology (2018). doi: 10.1007/s00418-018-1693-6

25. Chaikeeratisak, V. et al. Assembly of a nucleus-like structure during viral replication in bacteria. Science (80-.). (2017). doi: 10.1126/science.aal2130

26. Chaikeeratisak, V. et al. Viral Capsid Trafficking along Treadmilling Tubulin Filaments in Bacteria. Cell (2019). doi: 10.1016/j.cell.2019.05.032

27. Moniruzzaman, M., Martinez-Gutierrez, C. A., Weinheimer, A. R. & Aylward, F. O. Dynamic Genome Evolution and Blueprint of Complex Virocell Metabolism in Globally-Distributed Giant Viruses. bioRxiv (2019). doi: 10.1101/836445

28. Seitz, K. W. et al. Asgard archaea capable of anaerobic hydrocarbon cycling. Nat. Commun. (2019). doi: 10.1038/s41467-019-09364-x

29. Imachi, H. et al. Isolation of an archaeon at the prokaryote–eukaryote interface. Nature (2020). doi: 10.1038/s41586-019-1916-6

30. Seitz, K. W., Lazar, C. S., Hinrichs, K. U., Teske, A. P. & Baker, B. J. Genomic reconstruction of a novel, deeply branched sediment archaeal phylum with pathways for acetogenesis and sulfur reduction. ISME J. (2016). doi: 10.1038/ismej.2015.233

31. Spang, A. et al. Complex archaea that bridge the gap between prokaryotes and eukaryotes. Nature 521, 173–179 (2015).

32. Carradec, Q. et al. A global ocean atlas of eukaryotic genes. Nat. Commun. (2018). doi: 10.1038/s41467-017-02342-1

33. Eren, A. M. et al. Anvi’o: an advanced analysis and visualization platform for ‘omics data. PeerJ 3, e1319 (2015).

34. Delmont, T. O. et al. Nitrogen-fixing populations of Planctomycetes and Proteobacteria are abundant in surface ocean metagenomes. Nat. Microbiol. 3, (2018).

35. Guindon, S. et al. New algorithms and methods to estimate maximum-likelihood phylogenies: Assessing the performance of PhyML 3.0. Syst. Biol. 59, 307–321 (2010).

36. Hoang, D. T., Chernomor, O., Von Haeseler, A., Minh, B. Q. & Vinh, L. S. UFBoot2: Improving the ultrafast bootstrap approximation. Mol. Biol. Evol. (2018). doi: 10.1093/molbev/msx281

37. Katoh, K. & Standley, D. M. MAFFT multiple sequence alignment software version 7: Improvements in performance and usability. Mol. Biol. Evol. (2013). doi: 10.1093/molbev/mst010

38. Criscuolo, A. & Gribaldo, S. BMGE (Block Mapping and Gathering with Entropy): A new software for selection of phylogenetic informative regions from multiple sequence alignments. BMC Evol. Biol. 10, 210 (2010).

39. Nguyen, L. T., Schmidt, H. A., Von Haeseler, A. & Minh, B. Q. IQ-TREE: A fast and effective stochastic algorithm for estimating maximum-likelihood phylogenies. Mol. Biol. Evol. (2015). doi: 10.1093/molbev/msu300

40. Lemoine, F. et al. Renewing Felsenstein’s phylogenetic bootstrap in the era of big data. Nature (2018). doi: 10.1038/s41586-018-0043-0

41. Letunic, I. & Bork, P. Interactive Tree Of Life (iTOL): an online tool for phylogenetic tree display and annotation. Bioinformatics 23, 127–128 (2007).

42. Kelley, L. A., Mezulis, S., Yates, C. M., Wass, M. N. & Sternberg, M. J. E. The Phyre2 web portal for protein modeling, prediction and analysis. Nat. Protoc. (2015). doi: 10.1038/nprot.2015.053

